# The *C. elegans* female state: Decoupling the transcriptomic effects of aging and sperm-status

**DOI:** 10.1101/083113

**Authors:** David Angeles-Albores, Daniel H. W. Leighton, Tiffany Tsou, Tiffany H. Khaw, Igor Antoshechkin, Paul W. Sternberg

## Abstract

Understanding genome and gene function in a whole organism requires us to fully comprehend the life cycle and the physiology of the organism in question. Although *C. elegans* is traditionally thought of as a hermaphrodite, XX animals exhaust their sperm and become endogenous females after 3days of egg-laying. The molecular physiology of this state has not been as intensely studied as other parts of the life cycle, despite documented changes in behavior and metabolism that occur at this stage. To study the female state of *C. elegans*, we measured the transcriptomes of 1st day adult hermaphrodites; endogenous, 6th day adult females; and at the same time points, mutant *fog-2(lf)* worms that have a feminized germline phenotype. At these time points, we could separate the effects of biological aging from the transition into the female state. *fog-2(lf)* mutants partially phenocopy 6 day adult wild-type animals and exhibit fewer differentially expressed genes as they age throughout these 6 days. Therefore, *fog-2* is epistatic to age as assessed by this transcriptomic phenotype, which indicates that both factors act on sperm status to mediate entry into the female state. These changes are enriched in transcription factors canonically associated with neuronal development and differentiation. Our data provide a high-quality picture of the changes that happen in global gene expression throughout the period of early aging in the worm.

## Introduction

Transcriptome analysis by RNA-seq[1] has allowed for indepth analysis of gene expression changes between life stages and environmental conditions in many species [2,3].*Caenorhabditiselegans*, a genetic model nematode with extremely well defined and largely invariant development [4,5], has been subjected to extensive transcriptomic analysis across all stages of larval development [6–8] and many stages of embryonic development [7]. Although RNA-seq was used to develop transcriptional profiles of the mammalian aging process soon after its invention [9], few such studies have been conducted in *C. elegans* past the entrance into adulthood.

A distinct challenge to the study of aging transcriptomes in *C. elegans* is the hermaphroditic lifestyle of wild-type individuals of this species. Young adult hermaphrodites are capable of self-fertilization [10,11], and the resulting embryos will contribute RNA to whole-organism RNA extractions. Most previous attempts to study the *C. elegans* aging transcriptome have addressed the aging process only indirectly, or relied on the use of genetically or chemically sterilized animals to avoid this problem [7,12–17]. In addition, most of these studies obtained transcriptomes using microarrays, which are less accurate than RNA-seq, especially for low-expressed genes [18].

Here, we investigate what we argue is a distinct state in the *C. elegans* life cycle, the endogenous female state. Although *C. elegans* hermaphrodites emerge into adulthood already replete with sperm, after about 3 days of egg-laying the animals become sperm-depleted and can only reproduce by mating, This marks a transition into what we define as the endogenous female state. This state is behaviorally distinguished by increased male-mating success [19], which may be due to an increased attractiveness to males [20]. This increased attractiveness acts at least partially through production of volatile chemical cues [21]. These behavioral changes are also coincident with functional deterioration of the germline[22], muscle [23], intestine [24] and nervous system [25], changes traditionally attributed to the aging process [26].

To decouple the effects of aging and sperm-loss, we devised a two factor experiment. We examined wild-type XX animals at the beginning of adulthood (before worms contained embryos, referred to as 1st day adults) and after sperm depletion (6 days after the last molt, which we term 6th day adults). Second, we examined feminized XX animals that fail to produce sperm but are fully fertile if supplied sperm by mating with males (see Fig.1). We used *fog-2(lf)* mutants to obtain feminized animals. *fog-2* is involved in germ-cell sex determination in the hermaphrodite worm and is required for sperm production [27,28].

**Figure 1.**
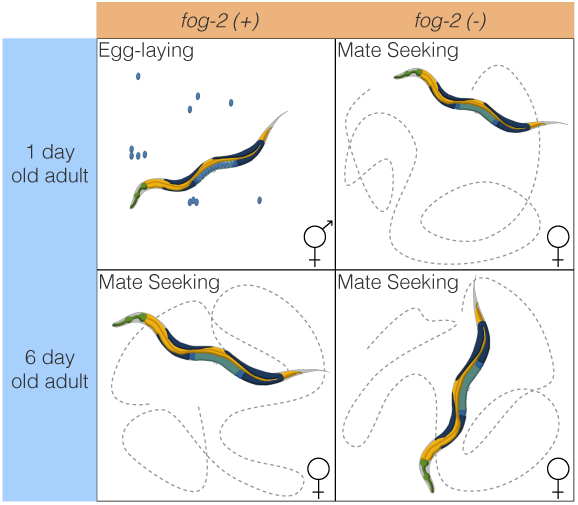
Experimental design to identify genes associated with sperm loss and with aging. Studying the wild-type worm alone would measure time- and sperm-related changes at the same time, without allowing us to separate these changes. Studying the wild-type worm and a *fog-2(lf)* mutant would enable us to measure sperm-related changes but not time-related changes. By mixing both designs, we can measure and separate both modules.

*C. elegans* defective in sperm formation will never transition into or out of a hermaphroditic stage. As time moves forward, these spermless worms only exhibit changes related to biological aging. We also reasoned that we might be able to identify gene expression changes due to different life histories: whereas hermaphrodites lay almost 300 eggs over three days, spermless females do not lay a single one. The different life histories could affect gene expression.

Here, we show that we can detect a transcriptional signature associated both with loss of hermaphroditic sperm and entrance into the endogenous female state. We can also detect changes associated specifically with biological aging. Loss of sperm leads to increases in the expression levels of transcription factors that are canonically associated with development and cellular differentiation and enriched in neuronal functions. Biological aging causes transcriptomic changes consisting of 5,592 genes in *C. elegans*. 4,552 of these changes occur in both genotypes we studied, indicating they do not depend on life history or genotype. To facilitate exploration of the data, we have generated a website where we have deposited additional graphics, as well as all of the code used to generate these analyses: https://wormlabcaltech.github.io/Angeles_Leighton_2016/.

## Materials and Methods

### Strains

Strains were grown at 20° C on NGM plates containing *E. coli* OP50. We used the laboratory *C. elegans* strain N2 as our wild-type strain [10]. We also used the N2 mutant strain JK574, which contains the *fog-2(q71)* allele, for our experiments.

### RNA extraction

Synchronized worms were grown to either young adulthood or the 6th day of adulthood prior to RNA extraction. Synchronization and aging were carried out according to protocols described previously [21]. 1,000-5,000 worms from each replicate were rinsed into a microcentrifuge tube in S basal (5.85g/L NaCl, 1g/L K_2_HPO_4_, 6g/L KH_2_PO_4_), and then spun down at 14,000rpm for 30s. The supernatant was removed and 1mL of TRIzol was added. Worms were lysed by vortexing for 30 s at room temperature and then 20 min at 4°. The TRIzol lysate was then spun down at 14,000rpm for 10 min at 4°C to allow removal of insoluble materials. Thereafter the Ambion TRIzol protocol was followed to finish the RNA extraction (MAN0001271 Rev. Date: 13 Dec 2012). 3 biological replicates were obtained for each genotype and each time point.

### RNA-Seq

RNA integrity was assessed using RNA 6000 Pico Kit for Bioanalyzer (Agilent Technologies #5067–1513) and mRNA was isolated using NEBNext Poly(A) mRNA Magnetic Isolation Module (New England Biolabs, NEB, #E7490). RNA-Seq libraries were constructed using NEBNext Ultra RNA Library Prep Kit for Illumina (NEB #E7530) following manufacturer’s instructions. Briefly, mRNA isolated from ~1*μ*g of total RNA was fragmented to the average size of 200nt by incubating at 94°C for 15 min in first strand buffer,cDNA was synthesized using random primers and ProtoScript II Reverse Transcriptase followed by second strand synthesis using Second Strand Synthesis Enzyme Mix (NEB). Resulting DNA fragments were end-repaired, dA tailed and ligated to NEBNext hairpin adaptors (NEB #E7335). After ligation, adaptors were converted to the ‘Y’ shape by treating with USER enzyme and DNA fragments were size selected using AgencourtAMPure XP beads (Beckman Coulter #A63880) to generate fragment sizes between 250 and 350 bp. Adaptor-ligated DNA was PCR amplified followed by AMPure XP bead clean up. Libraries were quantified with Qubit ds-DNA HS Kit (ThermoFisher Scientific #Q32854) and the size distribution was confirmed with High Sensitivity DNA Kit for Bioanalyzer (Agilent Technologies #5067–4626). Libraries were sequenced on Illumina HiSeq2500 in single read mode with the read length of 50nt following manufacturer’s instructions. Base calls were performed with RTA 1.13.48.0 followed by conversion to FASTQ with bcl2fastq 1.8.4.

## Statistical Analysis

### RNA-Seq Analysis

RNA-Seq alignment was performed using Kallisto[29] with 200 bootstraps. The commands used for read-alignment are in the S.I. file 1. Differential expression analysis was performed using Sleuth [30]. The following General Linear Model (GLM) was fit:

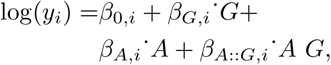

where *y_i_*are the TPM counts for the ith gene; *β*_0,*i*_ is the intercept for the ith gene, and *β_X,ί_* is the regression coefficient for variable *X* for the *i*th gene; *A* is a binary age variable indicating 1st day adult (0) or 6th day adult (1) and *G* is the genotype variable indicating wild-type (0) or *fog-2(lf)* (1); *β_A::G, i_* refers to the regression coefficient accounting for the interaction between the age and genotype variablesin the *i*^th^ gene. Genes were called significant if the FDR-adjusted q-value for any regression coefficient was less than 0.1. Our script for differential analysis is available on GitHub.

Regression coefficients and TPM counts were processed using Python 3.5 in a Jupyter Notebook [31]. Data analysis was performed using the Pandas, NumPy and SciPy libraries [32–34]. Graphics were created using the Matplotlib and Seaborn libraries [35,36]. Interactive graphics were generated using Bokeh[37].

Tissue, Phenotype and Gene Ontology Enrichment Analyses (TEA, PEA and GEA, respectively) were performed using the WormBase Enrichment Suite for Python [38,39].

## Data Availability

Strains are available from the *Caenorhabditis* Genetics Center. All of the data and scripts pertinent for this project except the raw reads can be found on our Github repository https://github.com/WormLabCaltech/Angeles_Leighton_2016. File S1 contains the list of genes that were altered in aging regardless of genotype. File S2 contains the list of genes and their associations with the *fog-2(lf)* phenotype. File S3 contains genes associated with the female state. Raw reads were deposited to the Sequence Read Archive under the accession code SUB2457229.

## Results and Discussion

### Decoupling time-dependent effects from sperm-status via general linear models

In order to decouple time-dependent effects from changes associated with loss of hermaphroditic sperm, we measured wild-type and *fog-2(lf)* adults at the 1st day adult stage (before visible embryos were present) and 6th day adult stage, when all wild-type hermaphrodites had laid all their eggs (see Fig1), but mortality was still low (< 10%) [41]. We obtained 16–19 million reads mappable to the *C. elegans* genome per biological replicate, which enabled us to identify 14,702 individual genes totalling 21,143 isoforms (see Figure2a).

**Figure 2.**
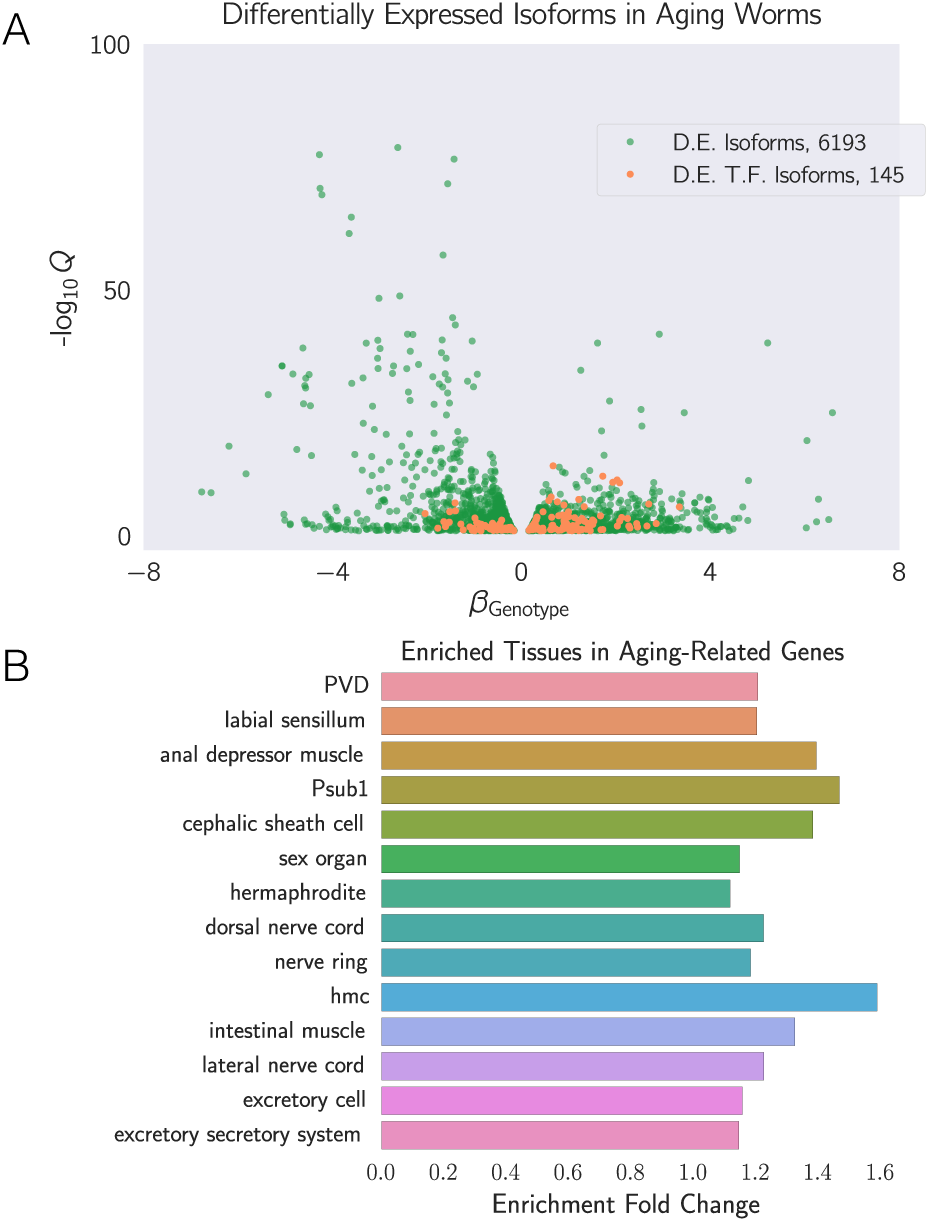
**A** We identified a common aging transcriptome between N2 and *fog-2(lf)* animals, consisting of 6,193 differentially expressed isoforms totaling 5,592 genes. The volcano plot is randomly down-sampled 30% for ease of viewing. Each point represents an individual isoform. *β*_Aging_ is the regression coefficient. Largei magnitudes of *β* indicate a larger log-fold change. The y-axis shows the negative logarithm of the q-values for each point. Green points are differentially expressed isoforms; orange points are differentially expressed isoforms of predicted transcription factor genes [40]. An interactive version of this graph can be found on our website **B** Tissue Enrichment Analysis [38] showed that genes associated with muscle tissues and the nervous system are enriched in aging-related genes. Only statistically significantly enriched tissues are shown. Enrichmenl Fold Change is defined as *Observed/Expected*. hmc stands for head mesodermal cell.

One way to analyze the data from this two-factor design is by pairwise comparison of the distinct states. However, such an analysis would not make full use of all the statistical power afforded by this experiment. Another method that makes full use of the information in our experiment is to perform a linear regression in 3 dimensions (2 independent variables, age and genotype, and 1 output). A linear regression with 1 parameter (age, for example) would fit a line between expression data for young and old animals. When a second parameter is added to the linear regression, said parameter can be visualized as altering the y-intercept, but not the slope, of the first line in question (see Fig.3a).

**Figure 3.**
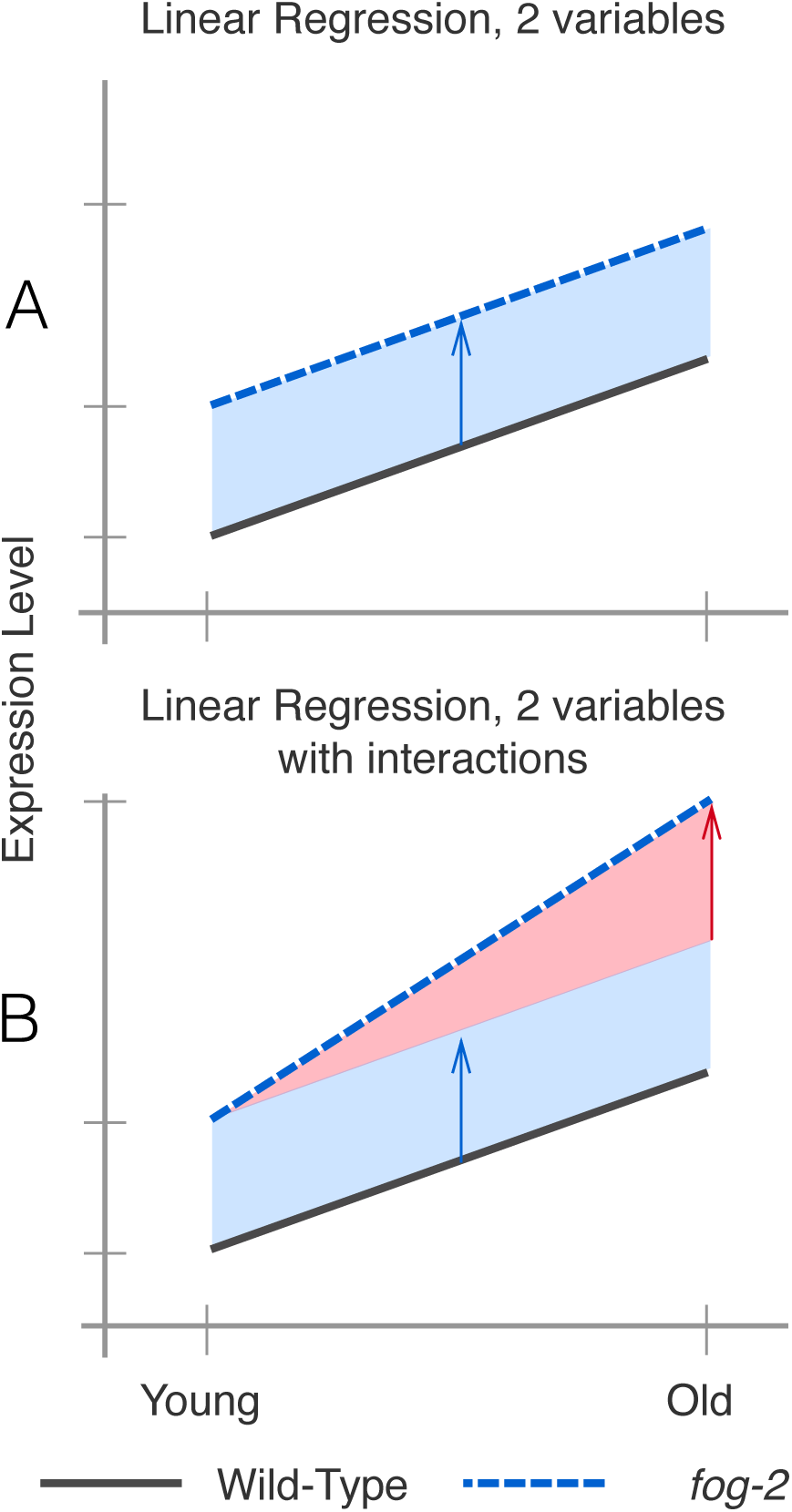
**A**. A linear regression with two variables, age and genotype. The expression level of a gene increases by the same amount as worms age regardless of genotype. However, *fog-2(lf)* has more mRNA than the wild-type at all stages (blue arrow). **B.** A linear regression with two variables and an interaction term. In this example, the expression level of this hypothetical gene is different between wild-type worms and *fog-2(lf)* (blue arrow). Although the expression level of this gene increases with age, the slope is different between wild-type and *fog-2(lf)*. The difference in the slope can be accounted for through an interaction coefficient (red arrow).

Although a simple linear model is oftentimes useful, sometimes it is not appropriate to assume that the two variables under study are entirely independent. For example, in our case, three out of the four timepoint-and-genotype combinations we studied did not have sperm, and sperm-status is associated with both the *fog-2(lf)* self-sterile phenotype and with biological age of the wild-type animal. One way to statistically model such correlation between variables is to add an interaction term to the linear regression. This interaction term allows extra flexibility in describing how changes occur between conditions. For example, suppose a given theoretical gene *X* has expression levels that increase in a *fog-2-dependent* manner, but also increases in an age-dependent manner. However, aged *fog-2(lf)* animals do not have expression levels of *X* that would be expected from adding the effect of the two perturbations; instead, the expression levels of *X* in this animal are considerably above what is expected. In this case, we could add a positive interaction coefficient to the model to explain the effect of genotype on the y-intercept as well as the slope (see Fig.3b). When the two perturbations are loss-of-function mutations, such interactions are epistatic interactions.

For these reasons, we used a linear generalized model (see Statistical Analysis) with interactions to identify a transcriptomic profile associated with the *fog-2(lf)* genotype independently of age, as well as a transcriptomic profile of *C. elegans* aging common to both genotypes. The change associated with each variable is referred as *β*; this number, although related to the natural logarithm of the fold change, is not equal to it. However, it is true that larger magnitudes of *β*indicate greater change. Thus, for each gene we performed a linear regression, and we evaluated the whether the *β*values associated with each coefficient were significantly different from 0 via a Wald test corrected for multiple hypothesis testing. A coefficient was considered to be significantly different from 0 if the q-value associated with it was less than 0.1.

### A quarter of all genes change expression between the 1st day of adulthood and the 6th day of adulthood in *C. elegans.*

We identified a transcriptomic signature consisting of 5,592 genes that were differentially expressed in 6th day adult animals of either genotype relative to 1st day adult animals (see SI file 2). This constitutes more than one quarter of the genes in *C. elegans.* Tissue Enrichment Analysis (TEA) [38] showed that nervous tissues including the ‘nerve ring’,‘dorsal nerve cord’,‘PVD’ and ‘labial sensillum’ were enriched in genes that become differentially expressed through aging. Likewise, certain muscle groups (‘anal depressor muscle’,‘intestinal muscle’) were enriched. (see Figure 2b). Gene Enrichment Analysis (GEA) [39] revealed that genes that were differentially expressed during the course of aging were enriched in terms involving respiration (‘respiratory chain’,‘oxoacid metabolic process’); translation (‘cytosolic large ribosomal subunit’); and nucleotide metabolism (‘purine nucleotide’,‘nucleoside phosphate’ and ‘ribose phosphate’ metabolic process). Phenotype Enrichment Analysis (PEA) [39] showed enrichment of phenotypes that affect the *C. elegans* gonad, including ‘gonad vesiculated’,‘gonad small’,‘oocytes lack nucleus’and ‘rachis narrow’.

To verify the quality of our dataset, we generated a list of 1,056 golden standard genes expected to be altered in 6th day adult worms using previous literature reports including downstream genes of *daf-12, daf-16*, and aging and lifespan extension datasets [12–16]. Out of 1,056 standard genes, we found 506 genes in our time-responsive dataset. This result was statistically significant with a p-value < 10^−38^.

Next, we used a published compendium [40] to search for known or predicted transcription factors. We found 145 transcription factors in the set of genes with differential expression in aging nematodes. We subjected this list of transcription factors to TEA to understand their expression patterns. 6 of these transcription factors were expressed in the ‘hermaphrodite specific neuron’ (HSN), a neuron physiologically relevant for egg-laying (*hlh-14, sem-4, ceh-20, egl-46, ceh-13, hlh-3*), which represented a statistically significant 2-fold enrichment of this tissue (*q* < 10^−1^). The term ‘head muscle’ was also overrepresented at twice the expected level (q < 10^−1^, 13 genes). Many of these transcription factors have been associated with developmental processess, and it is unclear why they would change expression in adult animals.

### The whole-organism *fog-2(lf)* transcriptome in *C. elegans*

We identified 1,881 genes associated with the *fog-2(lf)* genotype, including 60 transcription factors (see SI file 3). TEA showed that the terms ‘AB’,‘midbody’, ‘uterine muscle’,‘cephalic sheath cell’,‘anal depressor muscle’ and ‘PVD’ were enriched in this gene set. The terms ‘AB’ and ‘midbody’ likely reflect the impact of *fog-2(lf)* on the germline. Phenotype enrichment showed that only a single phenotype, ‘spindle orientation variant’ was enriched in the *fog-2(lf)*transcriptome (q < 10^−1^, 38 genes, 2-fold enrichment). Most genes annotated as ‘spindle orientation variant’ were slightly upregulated, and therefore are unlikely to uniquely reflect reduced germline proliferation. GO term enrichment was very similar to the aging gene set and reflected enrichment in annotations pertaining to translation and respiration. Unlike the aging gene set, the *fog-2(lf)*transcriptome was significantly enriched in ‘myofibril’ and ‘G-protein coupled receptor binding’ (q < 10^−1^). Enrichment of the term ‘G-protein coupled receptor binding’ was due to 14 genes: *cam-1, mom-2, dsh-1, spp-10, flp-6, flp-7, flp-9, flp-13, flp-14, flp-18, K02A11.4, nlp-12, nlp-13*, and *nlp-40. dsh-1, mom-2* and *cam-1* are members of the Wnt signaling pathway. Most of these genes’ expression levels were up-regulated, suggesting increased G-protein binding activity in *fog-2(lf)* mutants.

### The *fog-2(lf)* transcriptome overlaps significantly with the aging transcriptome

Of the 1,881 genes that we identified in the *fog-2(lf)* transcriptome, 1,040 genes were also identified in our aging set. Moreover, of these 1,040 genes, 905 genes changed in the same direction in response either aging or germline feminization. The overlap between these transcriptomes suggests an interplay between sperm-status and age. The nature of the interplay should be captured by the interaction coefficients in our model. There are four possibilities. First, the *fog-2(lf)* worms may have a fast-aging phenotype, in which case the interaction coefficients should match the sign of the aging coefficient. Second, the *fog-2(lf)* worms may have a slow-aging phenotype, in which case the interaction coefficients should have an interaction coefficient that is of opposite sign, but not greater in magnitude than the aging coefficient (if a gene increases in aging in a wild-type worm, it should still increase in a *fog-2(lf)* worm, albeit less). Third, the *fog-2(lf)* worms exhibit a rejuvenation phenotype. If this is the case, then these genes should have an interaction coefficient that is of opposite sign and greater magnitude than their aging coefficient, such that the change of these genes in *fog-2(lf)* mutant worms is reversed relative to the wild-type. Finally, if these genes are indicative of a female state, then these genes should not change with age in *fog-2(lf)* animals, since these animals do not exit this state during the course of the experiment. Moreover, because wild-type worms become female as they age, a further requirement for a transcriptomic signature of the female state is that aging coefficients for genes in this signature should have genotype coefficients of equal sign and magnitude. In other words, entrance into the female state should be not be path-dependent.

**Figure 4.**
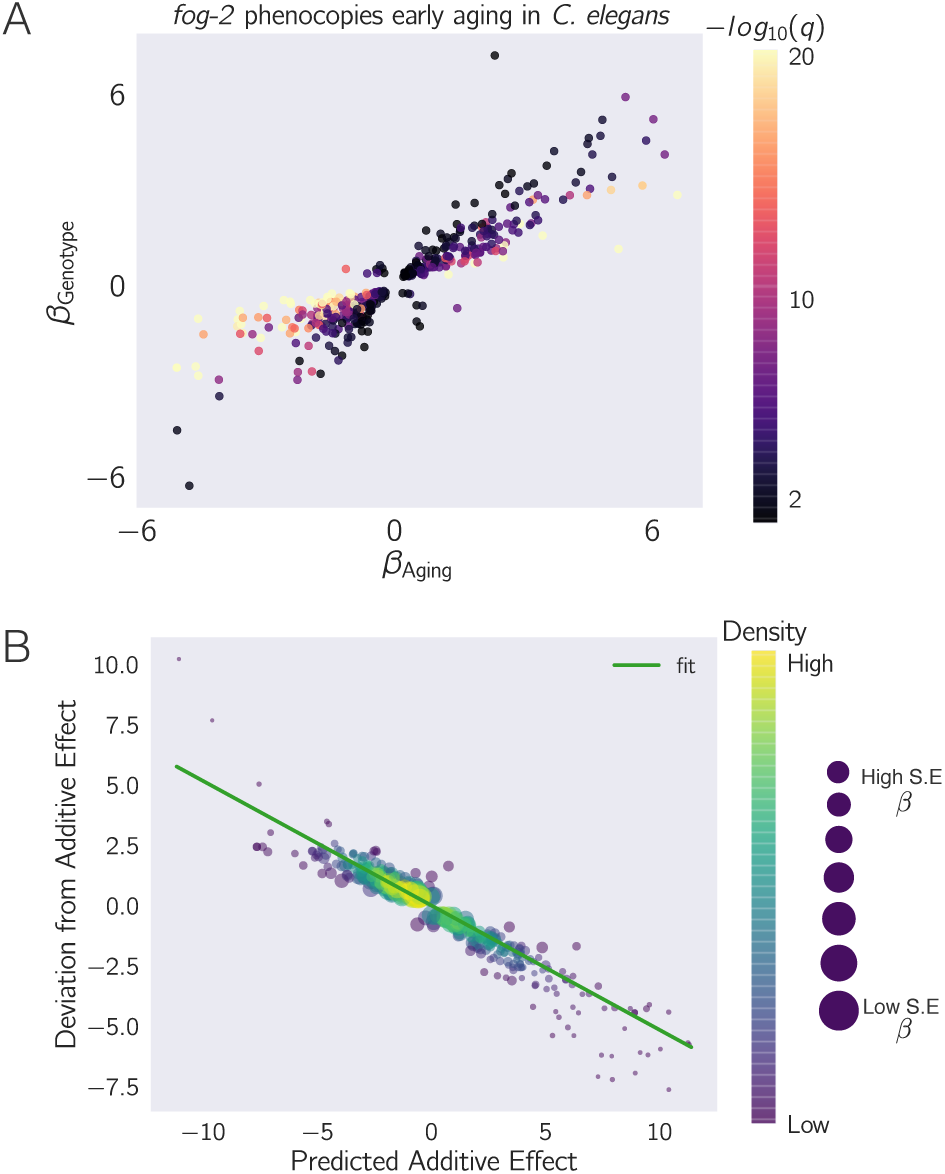
*fog-2(lf)* partially phenocopies early aging in *C. elegans*. The *β* in each axes is the regression coefficient from the GLM, and can be loosely interpreted as an estimator of the log-fold change. Feminization by loss of *fog-2(lf)* is associated with a transcriptomic phenotype involving 1,881 genes. 1,040/1,881 of these genes are also altered in wild-type worms as they progress from young adulthood to old adulthood, and 905 change in the same direction. However, progression from young to old adulthood in a *fog-2(lf)* background results in no change in the expression level of these genes. **A** We identified genes that change similarly during feminization and aging. The correlation between feminization and aging is almost 1:1. **B** Epistasis plot of aging versus feminization. Epistasis plots indicate whether two genes (or perturbations) act on the same pathway. When two effects act on the same pathway, this is reflected by a slope of –0.5. The measured slope was −0.51 ± 0.01.

To evaluate which of these possibilities was most likely, we selected the 1,040 genes that had aging, genotype and interaction coefficients significantly different from zero and we plotted their temporal coefficients against their genotype coefficients (see Fig. 4a). We observed that the aging coefficients were strongly predictive of the genotype coefficients. Most of these genes fell near the line *y* = *x*, suggesting that these genes define a female state. As a further test that these genes actually define a female state, we generated an epistasis plot using this gene set. We have previously used epistasis plots to measure transcriptome-wide epistasis between genes in a pathway [42]. Briefly, an epistasis plot plots the expected expression of a double perturbation under an additive model (null model) on the x-axis, and the deviation from this null model in the y-axis. In other words, we calculated the x-coordinates for each point by adding *β*^Genotype^ + *β*_Aging_, and the y-coordinates are equal to *β_interaction_*for each isoform. Previously we have shown that if two genes act in a linear pathway, an epistasis plot will generate a line with slope equal to −0.5. When we generated an epistasis plot and found the line of best fit, we observed a slope of –0.51 ± 0.01, which suggests that the *fog-2* gene and time are acting to generate a single transcriptomic phenotype along a single pathway. Overall, we identified 405 genes that increased in the same direction through age or mutation of the *fog-2(lf)* gene and that had an interaction coefficient of opposite sign to the aging or genotype coefficient (see SI file 4). Taken together, this information suggests that these 405 genes define a female state in *C. elegans*.

### Analysis of the Female State Transcriptome

To better understand the changes that happen after sperm loss, we performed tissue enrichment, phenotype enrichment and gene ontology enrichment analyses on the set of 405 genes that we associated with the female state. TEA showed no tissue enrichment using this gene-set. GEA showed that this gene list was enriched in constituents of the ribosomal subunits almost four times above background (*q* < 10^−5^, 17 genes). The enrichment of ribosomal constituents in this gene set in turn drives the enriched phenotypes: ‘avoids bacterial lawn’,‘diplotene absent during oogenesis’, ‘gonad vesiculated’,‘pachytene progression during oogenesis variant’, and ‘rachis narrow’. The expression of most of these ribosomal subunits is down-regulated in aged animals or in *fog-2(lf)* mutants.

**Figure 5.**
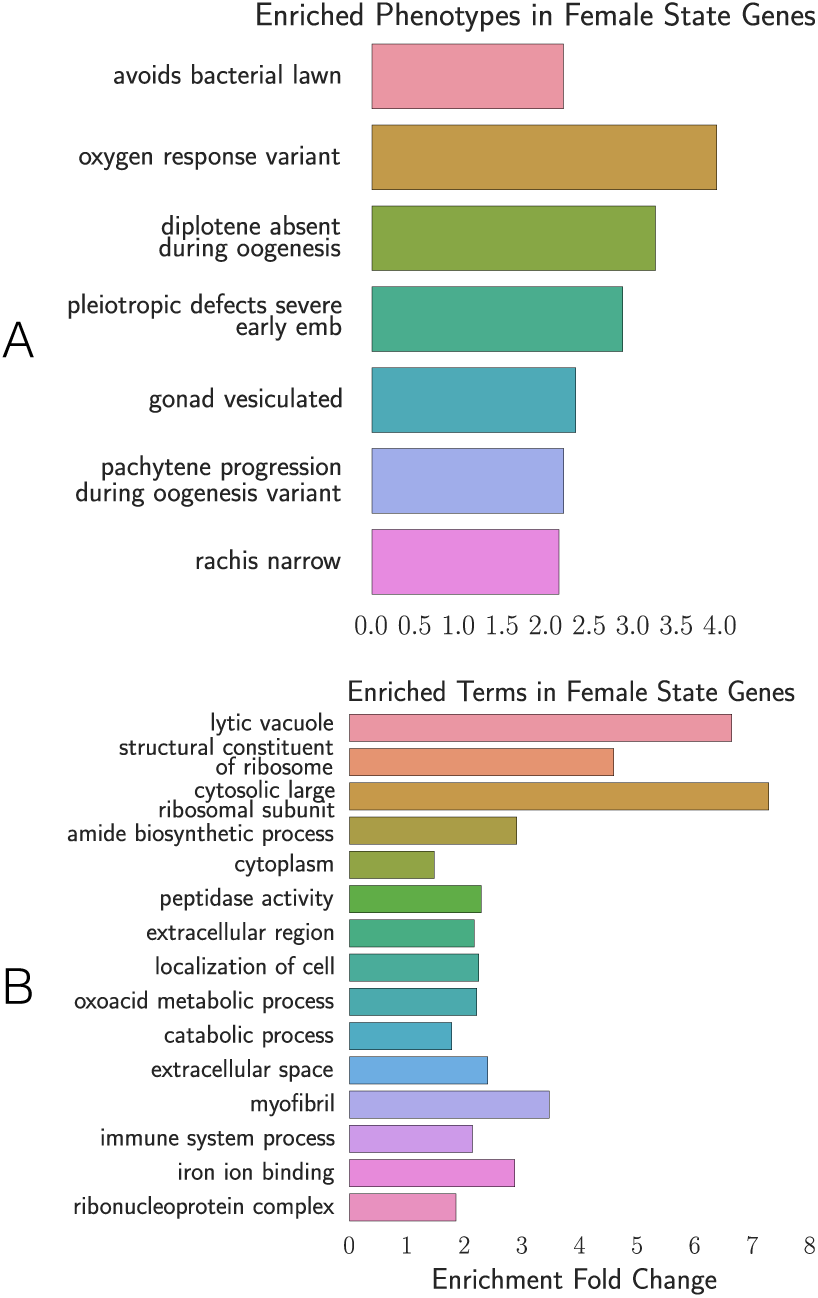
Phenotype and GO enrichment of genes involved in the female state. **A**. Phenotype Enrichment Analysis. **B.** Gene Ontology Enrichment Analysis. Most of the terms enriched in PEA reflect the abundance of ribosomal subunits present in this gene set.

## Discussion

### Defining an Early Aging Phenotype

Our experimental design enables us to decouple the effects of egg-laying from aging. As a result, we identified a set of almost 4,000 genes that are altered similarly between wild-type and *fog-2(lf)* mutants. Due to the read depth of our transcriptomic data (20 million reads) and the number of samples measured (3 biological replicates for 4 different life stages/genotypes), this dataset constitutes a high-quality description of the transcriptomic changes that occur in aging populations of *C. elegans*. Although our data only capture ~ 50% of the expression changes reported in earlier aging transcriptome literature, this disagreement can be explained by a difference in methodology; earlier publications typically addressed the aging of fertile wild-type hermaphrodites only indirectly, or queried aging animals at a much later stage of their life cycle.

### Measurement of a female state is enabled by linear models

We set out to study the self-fertilizing (hermaphroditic) to self-sterile (female) transition by comparing wild-type animals with *fog-2(lf)* mutants as they aged. Our computational approach enabled us to separate between two biological processes that are correlated within samples. Because of this intra-sample correlation, identifying this state via pairwise comparisons would not have been straightforward. Although it is a favored method amongst biologists, such pairwise comparisons suffer from a number of drawbacks. First, pairwise comparisons are unable to draw on the full statistical power available to an experiment because they discard almost all information except the samples being compared. Second, pairwise comparisons require a researcher to define *a priori* which comparisons are informative. For experiments with many variables, the number of pairwise combinations is explosively large. Indeed, even for this two-factor experiment, there are 6 possible pairwise comparisons. On the other hand, by specifying a linear regression model, each gene can be summarized with three variables, each of which can be analyzed and understood without the need to resort to further pairwise combinations.

Our explorations have shown that the loss of *fog-2(lf)* partially phenocopies the transcriptional events that occur naturally as *C. elegans* ages from the 1st day of adulthood to the 6th day of adulthood. Moreover, epistasis analysis of these perturbations suggest that they act on the same pathway, namely sperm generation and depletion (see Fig. 6). Sperm generation promotes a non-female states, whereas sperm depletion causes entry into the female state. Given the enrichment of neuronal transcription factors that are associated with sperm loss in our dataset, we believe this dataset should contain some of the transcriptomic modules that are involved in these pheromone production and behavioral pathways, although we have been unable to find these genes. Currently, we cannot judge how many of the changes induced by loss of hermaphroditic sperm are developmental (i.e., irreversible), and how many can be rescued by mating to a male. While an entertaining thought experiment, establishing whether these transcriptomic changes can be rescued by males is a daunting experimental task, given that the timescales for physiologic changes could reasonably be the same as the timescale of onset of embryonic transcription. All in all, our research supports the idea that wide-ranging transcriptomic effects of aging in various tissues can be observed well before onset of mortality, and that *C. elegans* continues to develop as it enters a new state of its life cycle.

**Figure 6.**
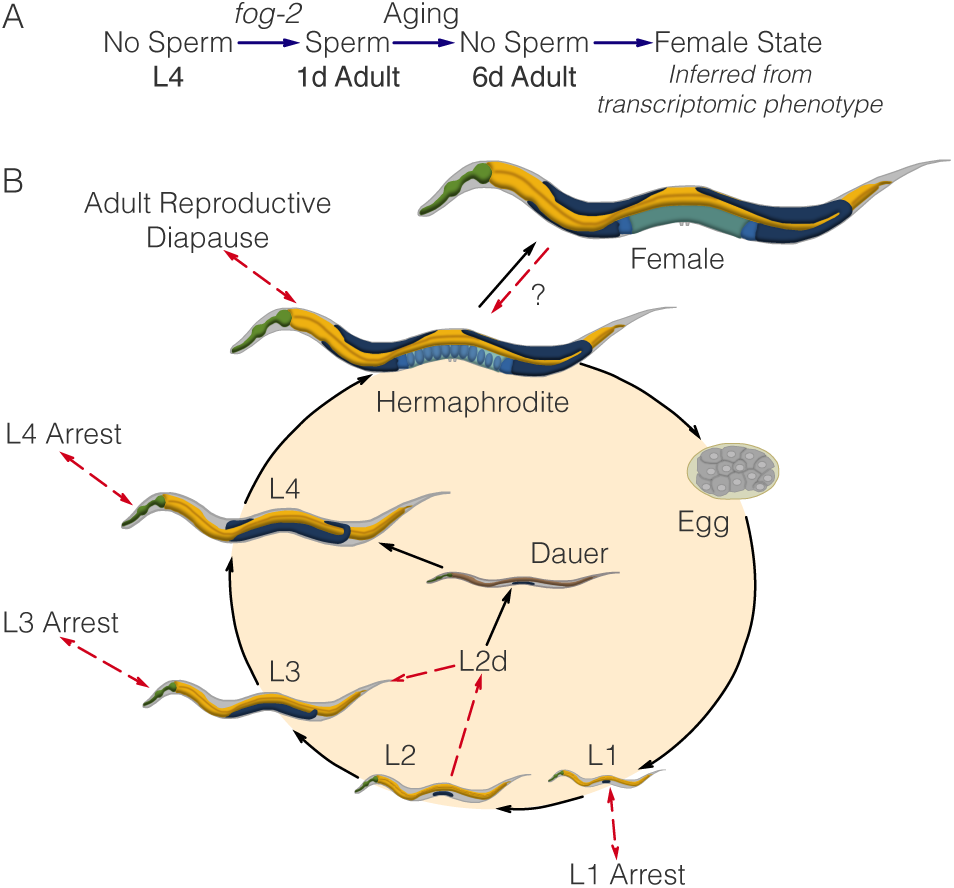
**A**. A substrate model showing how *fog-2* promotes sperm generation, whereas aging promotes sperm depletion, leading to entry to the female state. Such a model can explain why *fog-2* and aging appear epistatic to each other. **B**. The complete *C. elegans* life cycle. Recognized stages of *C. elegans* are marked by black arrows. States are marked by red arrows to emphasize that at the end of a state, the worm returns to the developmental timepoint it was at before entering the state. The L2d state is an exception. It is the only stage that does not return to the same developmental timepoint; rather, the L2d state is a permissive state that allows entry into either dauer or the L3 stage. We have presented evidence of a female state in *C. elegans*. At this point, it is unclear whether the difference between hermaphrodites and females is reversible by males. Therefore, it remains unclear whether it is a stage or a true state.

### The *C. elegans* life cycle, life stages and life states

*C. elegans* has a complicated life cycle, with two alternative developmental pathways that have multiple stages (larval development and dauer development), followed by reproductive adulthood. In addition to its developmental stages, researchers have recognized that *C. elegans* has numerous life states that it can enter into when given instructive environmental cues. One such state is the L1 arrest state, where development ceases entirely upon starvation [43]. More recently, researchers have described additional diapause states that the worm can access at the L3, L4 and young adult stages under conditions of low food [44–46]. Not all states of *C. elegans* are arrested, however (see Fig. 6). For example, the L2d state is induced by crowded and nutrient poor conditions [47]. While within this state, the worm is capable of entry into either dauer or the L3 larval stage, depending on environmental conditions. Thus, the L2d state is a permissive state, and marks the point at which the nematode development is committed to a single developmental pathway.

Identification of the *C. elegans* life states has often been performed by morphological studies (as in the course of L4 arrest or L2d) or via timecourses (L1 arrest). However, not all states may be visually identifiable, or even if they are, the morphological changes may be very subtle, making positive identification difficult. However, the detailed information afforded by a transcriptome should in theory provide sufficient information to definitively identify a state, since transcriptomic information underlies morphology. Moreover, transcriptomics can provide an informative description into the physiology of complex metazoan life state's via measurements of global gene expression. By identifying differentially expressed genes and using ontology enrichment analyses to identify gene functions, sites of expression or phenotypes that are enriched in a given gene set, researchers can obtain a clearer picture of the changes that occur in the worm in a less biased manner than by identifying gross morphological changes. RNA-seq is emerging as a powerful technology that has been used successfully in the past as a qualitative tool for target acquisition. More recent work has successfully used RNA-seq to establish genetic interactions between genes [48,49]. In this work, we have shown that whole-organism RNA-seq data can also be analyzed via a similar formalism to successfully identify internal states in a multi-cellular organism.

## Acknowledgments

We thank the *Caenorhabditis* Genetics Center for providing worm strains. This work would not be possible without the central repository of *C. elegans*information generated by WormBase, without which mining the genetic data would not have been possible. DHWL was supported by a National Institutes of Health US Public Health Service Training Grant (T32GM07616). This research was supported by the Howard Hughes Medical Institute, for which PWS is an investigator.

## Author Contributions

DA, DHWL and PWS designed all experiments. DHWL and THK collected RNA for library preparation. IA generated libraries and performed sequencing. DA performed all bioinformatics and statistical analyses. DA, TT and DHWL performed all screens. DA, DHWL and PWS wrote the paper.

